# Testicular mosaicism in non-mosaic postpubertal Klinefelter patients with focal spermatogenesis and in non-mosaic prepubertal Klinefelter boys

**DOI:** 10.1101/2023.12.19.572320

**Authors:** Semir Gül, Veerle Vloeberghs, Inge Gies, Ellen Goossens

**Author notes:** **Corresponding Author:** Semir Gül, Faculty of Medicine, Histology and Embryology Department, Malatya Turgut Özal University, Battalgazi, 44210 Malatya, Turkey.

## Abstract

The aim of the study is to investigate testicular mosaicism in non-mosaic postpubertal Klinefelter Syndrome patients and in non-mosaic prepubertal Klinefelter boys Testes of the males with non-mosaic Klinefelter Syndrome at different developmental stages were used.

Immunohistochemical and fluorescent in situ hybridization analyses were applied for X chromosome ploidy in testis-specific cells in testicular biopsy samples from non-mosaic Klinefelter Syndrome patients.

According to our findings, all analyzed spermatogonia in both postpubertal and prepubertal non-mosaic Klinefelter Syndrome patients have a 46,XY karyotype. However, while the Sertoli cells surrounding spermatogonia in postpubertal samples also have a 46,XY karyotype, the Sertoli cells surrounding spermatogonia in prepubertal samples have a 47,XXY karyotype. Peritubular myoid cells and Leydig cells may also have mosaicism in both postpubertal patients and prepubertal boys.

In conclusion, we confirmed in situ using cell-specific markers that testicular mosaicism exists in non-mosaic Klinefelter Syndrome patients. Therefore, we hypothesize that focal spermatogenesis seen in some postpubertal Klinefelter Syndrome patients originates from euploid spermatogonia and Sertoli cells. Additionally, our findings suggest that only spermatogonia that have lost their X chromosome can survive. Furthermore, our data suggest that spermatogonia lose the extra X chromosome during fetal or neonatal life, while Sertoli cells lose it around puberty. These findings will lay the groundwork for new studies on exactly when and by which mechanism an extra X chromosome is lost in spermatogonia and Sertoli cells.

## INTRODUCTION

Klinefelter syndrome (KS) is a genetic aneuploid condition with an incidence of 1 to 2 per 1000 live male births characterized by a karyotype in which males have one or more additional copies of the X chromosome (47,XXY; 48,XXXY; 49,XXXXY) (1, 2). . About 80 to 90 percent of men diagnosed with KS are non-mosaic, meaning their cells all have the same karyotype. The remaining 10 to 20 percent are mosaic patients of whom some cells have a karyotype of 47,XXY (or a higher-grade aneuploidy), while other cells have a 46,XY karyotype (2–4). The presence of mosaicism is usually determined by karyotyping about 20-50 lymphocytes (5, 6).

KS is usually diagnosed in adulthood when men seek help from a reproductive medicine center. The primary features of KS in terms of reproductive biology are small, poorly functioning testicles, impaired reproductive metabolism, and sexual dysfunction resulting in infertility or reduced fertility (7–12). Infertility in KS patients is mainly due to the loss of germ cells, which accounts for 90% of azoospermia cases in non-mosaic KS patients (13). While spermatogenesis occurs in most of the seminiferous tubules in healthy fertile individuals, it rarely occurs in individuals with non-mosaic KS (13, 14). If focal spermatogenesis is present, KS patients are able to have genetically own children by medically assisted reproduction: testicular sperm extraction (TESE) in combination with intracytoplasmic sperm injection (ICSI) (15).

Previous studies have shown that analyzing testicular biopsy specimens from KS patients may result in three different findings: (1) samples with spermatozoa (16); (2) samples with only spermatogonia; and (3) samples lacking germ cells (17). Why some patients still have active spermatogenesis, while others have only spermatogonia or no germ cells remains unclear. It has been hypothesized that focal spermatogenesis is related to testicular mosaicism. Sciurano et al. claimed that, although Sertoli cells showed aneuploidy, spermatogonia in areas with focal spermatogenesis were mosaic and that spermatogonia with normal karyotype were the source of focal spermatogenesis (14). Similarly, Blanco et al. and Bergere et al. reported testicular mosaicism in non-mosaic KS patients with focal spermatogenesis (18, 19). However, the karyotype of the testis-specific cells in focal spermatogenic regions was not shown *in situ*. Therefore, it is still unclear what the karyotype of the cells that originate focal spermatogenesis is. In order to answer this question, a spatial and cell-specific analysis of the X chromosome ploidy should be performed.

In females, one of the X chromosomes is active (Xa), while the other one is repressed by various molecular mechanisms to form an inactive X (Xi) chromosome, called the Barr Body (20). Also in KS men, the extra X chromosome(s) are inactivated (21). X-inactive specific transcript (XIST) mRNA is the major molecule for the inactivation of X chromosomes (22, 23), but other modifications are also involved in the formation of Xi, e.g. histone-3-lysine-27-tri-methylation (H3K27me3) (24).. Although H3K27me3 is a genome-wide chromatin marker (25), since it accumulates in excessive amounts on the Xi chromosome, this chromosome is tightly packed. Immunohistochemical staining for H3K27me3 can be used to visualize the Xi chromosome, showing a distinct punctate pattern, usually at the nuclear periphery (26–28) (Figure 1). Schaefer et al. reported that H3K27me3 immunohistochemical staining detects the Xi with high sensitivity and specificity, and can be used to exhibit the sex of cells in tissues (28). Therefore, staining for H3K27me3 in combination with cell-specific markers enables cell-specific analysis of X chromosome ploidy. Detection of the punctate pattern of H3K27me3 means that there is an Xi chromosome (and thus at least two X chromosomes) present in the cell (28).

**Figure 1.**
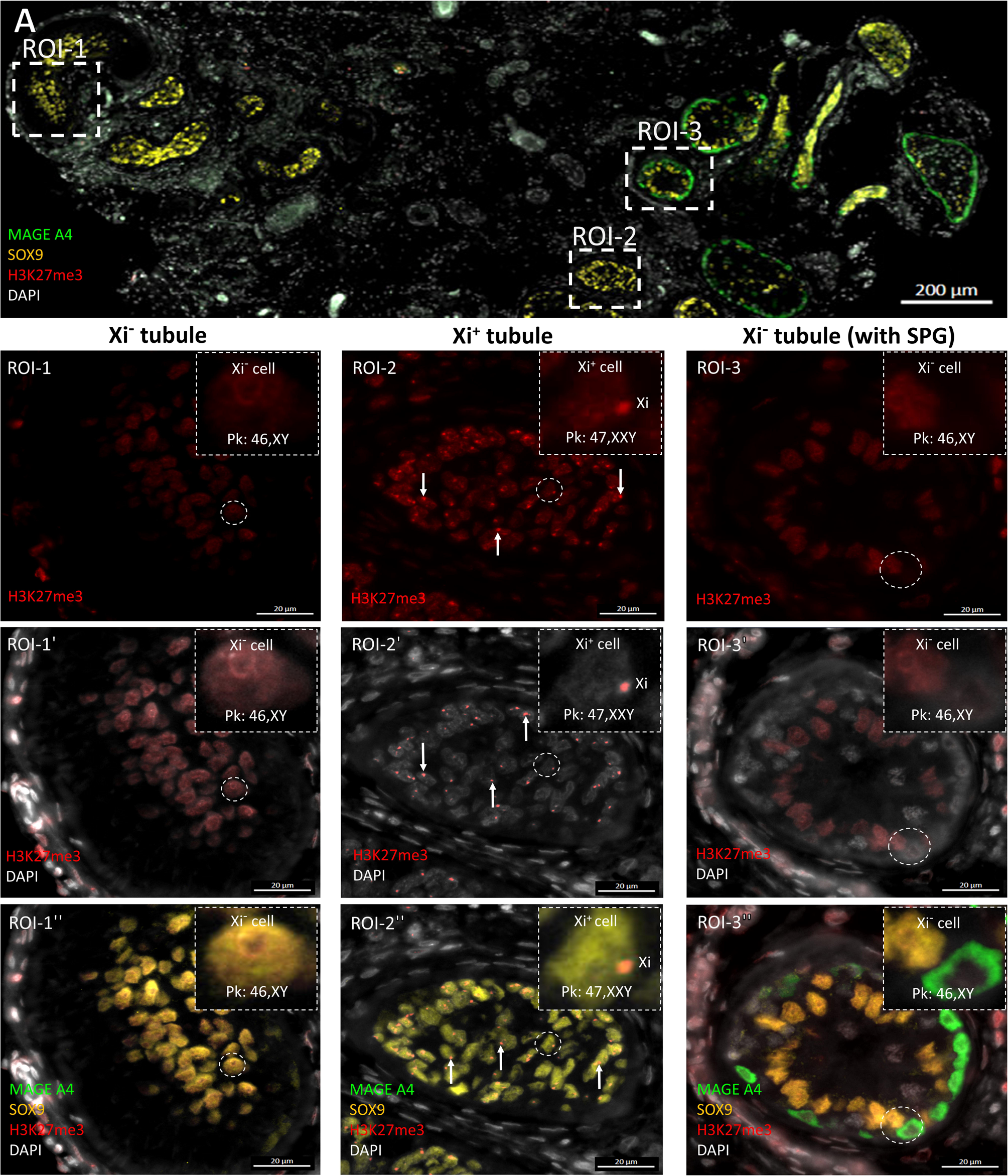
Classification of seminiferous tubules in KS testis. **A:** General view of testis section containing both SCO tubules and tubules with spermatogonia. **ROI-1, ROI-1’, ROI-1”:** A seminiferous tubule classified as Xi^-^ tubule. All Sertoli cells in the tubule are Xi^-^. The predicted karyotype is 46,XY. One of the Sertoli cells (encircled) is magnified in insets. **ROI-2, ROI-2’, ROI-2”:** A seminiferous tubule classified as Xi^+^ tubule. Almost all of the Sertoli cells in the tubule are Xi^+^. The predicted karyotype is 47,XXY. White arrows show Xi in Sertoli cells. One of the Sertoli cells (encircled) is magnified in insets. **ROI-3, ROI-3’, ROI-3”:** An Xi^-^ tubule with spermatogonia. All of the spermatogonia and Sertoli cells in the tubule are Xi^-^. The predicted karyotype for both is 46,XY. A Sertoli cell and a spermatogonium adjacent to each other (encircled) are magnified in insets. **Note:** DAPI was not applied in IF staining because DAPI application prevents the binding of FISH probes, so the pale DAPI colour (white) in IF figures was obtained by autofluorescence of nuclei using the DAPI channel. ROI: region of interest, SPG: spermatogonium, Xi^-^ cell: a cell without Xi, Xi^+^ cell: a cell with Xi, Xi^-^ tubule: seminiferous tubule with all Xi^-^ Sertoli cells, Xi^+^ tubule: seminiferous tubule with Xi^+^ (and Xi^-^) Sertoli cells.

Since it remains unclear whether there is mosaicism in germ cells and testis-specific somatic cells in non-mosaic KS patients, the aim of this study is to reveal the karyotypes of spermatogonia, Sertoli cells, peritubular myoid cells (PTMC) and Leydig cells in non-mosaic postpubertal KS patients with focal spermatogenesis and in non-mosaic prepubertal KS boys with presence of germ cells.

## MATERIALS AND METHODS

### Tissue samples

The use of tissue was approved by the ethics committee of the University Hospital in Brussels (UZ Brussel), Belgium (2015/121; BUN 143201524312). A total of 22 patients (17 postpubertal and 5 prepubertal) who were non-mosaic according to lymphocyte karyotype analysis, were included in the study (Supplementary table 1). Euploid female colon tissue (presence of Xi; XX control) and euploid testicular tissue (absence of Xi; XY control) were used as control samples (Supplementary table 1).

### Immunofluorescent staining

Tissue fixation, processing, sectioning (4 µm thickness), and deparaffinization were done as previously described (29). The protocol can be found in supplementary file 1.For each sample, one section was stained with melanoma-associated antigen 4 (MAGE A4) for spermatogonia (30), sex determining region Y-box 9 (SOX9) for Sertoli cells (31) and H3K27me3 for Xi (26–28, 32); the other section was stained with cytochrome P450 17A1 (CYP17A1) for Leydig cells (33), actin alpha 2 (ACTA2) for PTMCs (34) and H3K27me3 for Xi. The primary antibodies (Supplementary table 2) were incubated at 4°C overnight. For the negative control, only PBS with 1% normal donkey serum without the primary antibody was added onto the sections. Slides were washed in PBS (3x10 min) and secondary antibodies (Supplementary table 2) were incubated at RT for 1 hour. After the washing and mounting with ProLong^TM^ Gold antifade mounting medium (P36934, Invitrogen, Massachusetts, USA), slides were scanned by Zeiss Axio Scan.Z1 under 40x objective and images were saved.

### Fluorescence *in situ* hybridization (FISH)

FISH analysis was performed to assess the number of X chromosomes in a cell and validate the immunofluorescence results. Therefore, FISH was applied to the same sections following immunofluorescence staining. FISH staining was performed only on samples with spermatogonia (P1-P8, P18-P20). The coverslips were removed from the slides by immersing the slides in PBS for 15-30 minutes. Then, the FISH protocol was applied with probes for chromosome 18, X and Y (Z-2163-200, ZytoVision) according to manufacturer’s instructions (Z-2028-20, ZytoLight FISH-Tissue implementation Kit) (Supplementary file 1). Stained sections were scanned by Zeiss Axio Scan.Z1 under 40x objective and images were saved.

### Microscopic analysis and karyotyping

Images were analyzed by the software ZEISS ZEN 2.6 (blue edition). Cell-specific Xi analysis was done on immunofluorescent images and the results were confirmed on the FISH images. For each cell type, cells with a prominent H3K27me3 stained dot at the nuclear periphery were counted as Xi^+^; cells without H3K27me3 stained dot were counted as Xi^-^ (Figure 1) (28).

Seminiferous tubules were classified into two groups: Xi^+^ tubules (tubules containing at least one Sertoli cell with Xi) and Xi^-^ tubules (tubules containing exclusively Sertoli cells without Xi) (Figure 1). All seminiferous tubules found in sections were analyzed for spermatogonia and Sertoli cells.

For PTMC analysis, ten randomly selected seminiferous tubules were used. If less than ten tubules were present in the section, all tubules were evaluated. Leydig cell analyses were performed on 3-10 randomly selected areas (0.028±0.013 mm^2^ per section). As only one prepubertal sample was positive for CYP17A1, prepubertal samples were not evaluated for Leydig cells.

Per sample, the mean of Xi^+^ and Xi^-^ cells was calculated for each cell type, as well as the percentage of Xi^+^ cells. Xi^+^ cells were assessed as having the 47,XXY karyotype, while Xi^-^ cells were assessed as having the 46,XY karyotype (28). As a positive control for the presence of Xi, muscle cells in ten randomly selected areas in three sections taken at different depths from the female colon sample were analyzed.

X chromosome numbers of spermatogonia and Sertoli cells were confirmed by FISH. FISH signals for chromosomes 18, X and Y could only be counted in histologically intact sections of three postpubertal patients since FISH caused morphological distortion of the sections and shedding of cells from the slide. A total of 31 spermatogonia (4, 18, and 9 spermatogonia in P2, P8, and P9 respectively), 143 Sertoli cells (26 Sertoli cells in P2 and 117 Sertoli cells in P9) in Xi^-^ tubules and 233 Sertoli cells (56 Seroli cells in P2 and 177 Sertoli cells in P9) in Xi^+^ tubules were examined for FISH analysis.

### Statistical analysis

FISH results are shown as mean ±SD. The analysis of FISH signals was done with SPSS25. The Kolmogorov Smirnov Test was used to check whether the data included in the study fit the normal distribution. Since the variables did not have a normal distribution, the analysis was continued with non-parametric tests. The significance level (p) for comparison tests was set at 0.05. The Kruskal Wallis test analysis was performed for comparisons in multiple independent groups. As a result of the analysis, Bonferroni corrected p value was calculated.

## RESULTS

Detailed information about the patients and biopsy samples is given in Supplementary table 1. All patients were non-mosaic 47,XXY, according to lymphocyte karyotyping.

### Spermatogonia analysis

In 10 out of 17 postpubertal patients, sperm were obtained by TESE. However, in 5 of them (P9, P10, P13, P14, P15) spermatogonia could not be found in the cross-sections. In 3 postpubertal patients in which sperm retrieval by TESE was not successful (P5, P6, P8), tubules with spermatogonia were found (Supplementary table 1). Spermatogonia were detected in 41 of 118 Xi^-^ tubules of 8 postpubertal samples (P1-P8), and in 182 of 1297 Xi^+^ tubules of 3 prepubertal samples (P18-P20). Exceptionally, there were two Xi^+^ tubules with spermatogonia in P2 (supplementary figure 1). In these two tubules, spermatogonia were Xi^-^, the tubular structure and cells were disorganized, the tubular lumen was unclear, and there was no ongoing spermatogenesis.

In total, 826 spermatogonia in postpubertal samples, whether or not being in tubules with ongoing spermatogenesis, and 563 spermatogonia in prepubertal samples were analyzed. All spermatogonia, without exception, were Xi^-^. (Table 1, Figure 2-3) similar to the 46,XY control sample (Figure 2). According to FISH results, there were two signals for chromosome 18 and only one signal was found for X and Y chromosomes in spermatogonia nuclei (Figure 2-3) similar to the 46,XY control sample (Figure 2). In addition, the incidence of X and Y chromosome signals for spermatogonia in Xi^-^ tubules were similar, while more chromosome 18 signals were detected (p<0.001) (Figure 2). Therefore, both the immunofluorescence and FISH results suggest that the karyotype of all spermatogonia in both postpubertal and prepubertal samples is 46,XY.

**Figure 2.**
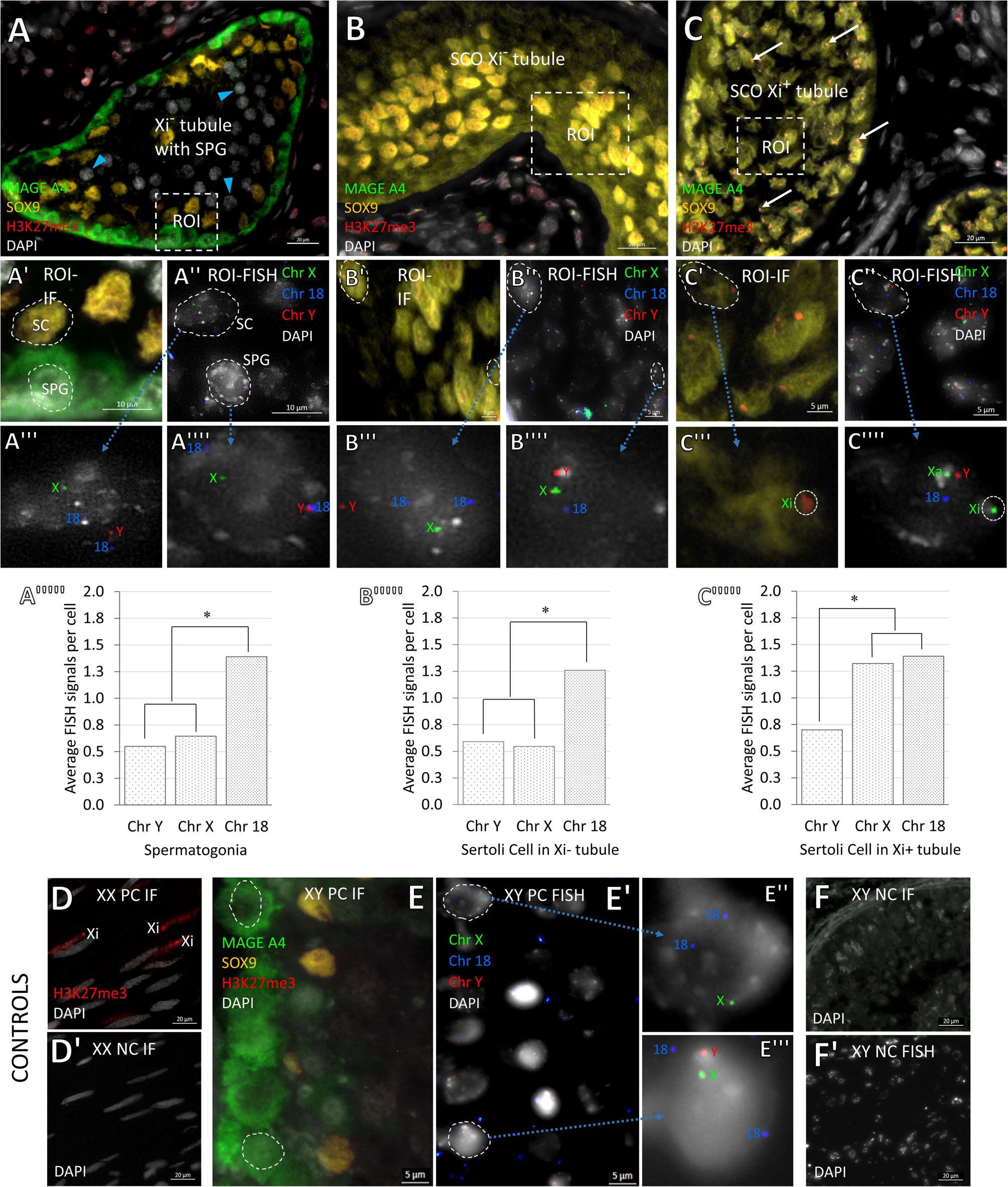
Xi and FISH analysis for spermatogonia and Sertoli cells in postpubertal sample. **A:** All spermatogonia and Sertoli cells are Xi^-^ in an Xi^-^ tubule with spermatogonia. Arrow heads show meiotic germ cells. **A’ and A”:** IF and FISH stain of ROI in A, respectively. One spermatogonium and one Sertoli cell are encircled. Both spermatogonia and Sertoli cells are Xi^-^ and have only one chr X signal. **A”’:** Magnified Sertoli cell in A” with two chr 18, one chr X, and one chr Y signals. **A””:** Magnified spermatogonium in A” with two chr 18, one chr X, and one chr Y signals. **A’”:** In spermatogonia, the incidence of X and Y chromosome FISH signals was similar, but about half as much as chromosome 18 signals. *p<0.05. **B:** A SCO tubule classified as Xi^-^ tubule. All Sertoli cells are Xi^-^. **B’ and B”:** IF and FISH stain of ROI in image B, respectively. Two Sertoli cells are encircled. All Sertoli cells are Xi^-^ in B’ and have one chr X signal in B”. **B”’:** A Sertoli cell magnified from B” with two chr 18, one chr X, and one chr Y signals. **B””:** Another Sertoli cell magnified from B” with one chr 18, one chr X, and one chr Y signals. One of the chr 18 signals was not observed because it did not coincide with the section plane. **B””’:** In Sertoli cells in Xi^-^ tubules, the incidence of X and Y chromosome signals was similar, but about half as much as chromosome 18 signals. *p<0.05. **C:** A SCO tubule classified as Xi^+^ tubule. The majority of the Sertoli cells are Xi^+^ (arrows). **C’ and C”:** IF and FISH stain of ROI in image C, respectively. One Sertoli cell is encircled. Most Sertoli cells are Xi^+^ in C’ and have two chr X signals in C”. **C”’ and C””:** IF and FISH stain of the same Sertoli cell. Encircled Xi in C”’ overlaps with one (encircled-Xi) of two chr X signals in C””. So, the second green signal is the active X (Xa). One chr 18 and one chr Y signal are also seen in C””. One of the chr 18 signal was not observed because it did not coincide with the section plane. **C””’:** In Sertoli cells in the Xi^+^ tubules, the incidence of chromosome 18 and X signals was similar, but approximately twice that of Y chromosome signal. *p<0.05. **D:** Muscle cells of colon tissue from a female as positive control for Xi analysis. **D’:** Negative control of female colon tissue for IF. **E:** 46,XY control testis tissue. All spermatogonia and Sertoli cells are Xi^-^. **E’:** FISH stain of the same area in E. **E” and E”’:** Two magnified spermatogonia, encircled in image E**’**, with two chr 18, one X, and one chr Y signal (in the bottom). Chr Y signal was not observed in the upper cell because it did not coincide with the section plane. **F:** Negative control for IF stain on testis tissue from the same XY control sample. **F’:** Negative control for FISH stain on testis from the same XY control sample. **Note:** DAPI was not applied in IF staining because DAPI application prevents the binding of FISH probes, so the pale DAPI colour (white) in IF figures was obtained by autofluorescence of nuclei using the DAPI channel. ROI: region of interest, SCO: Sertoli cell-only, SPG: spermatogonium.

**Figure 3.**
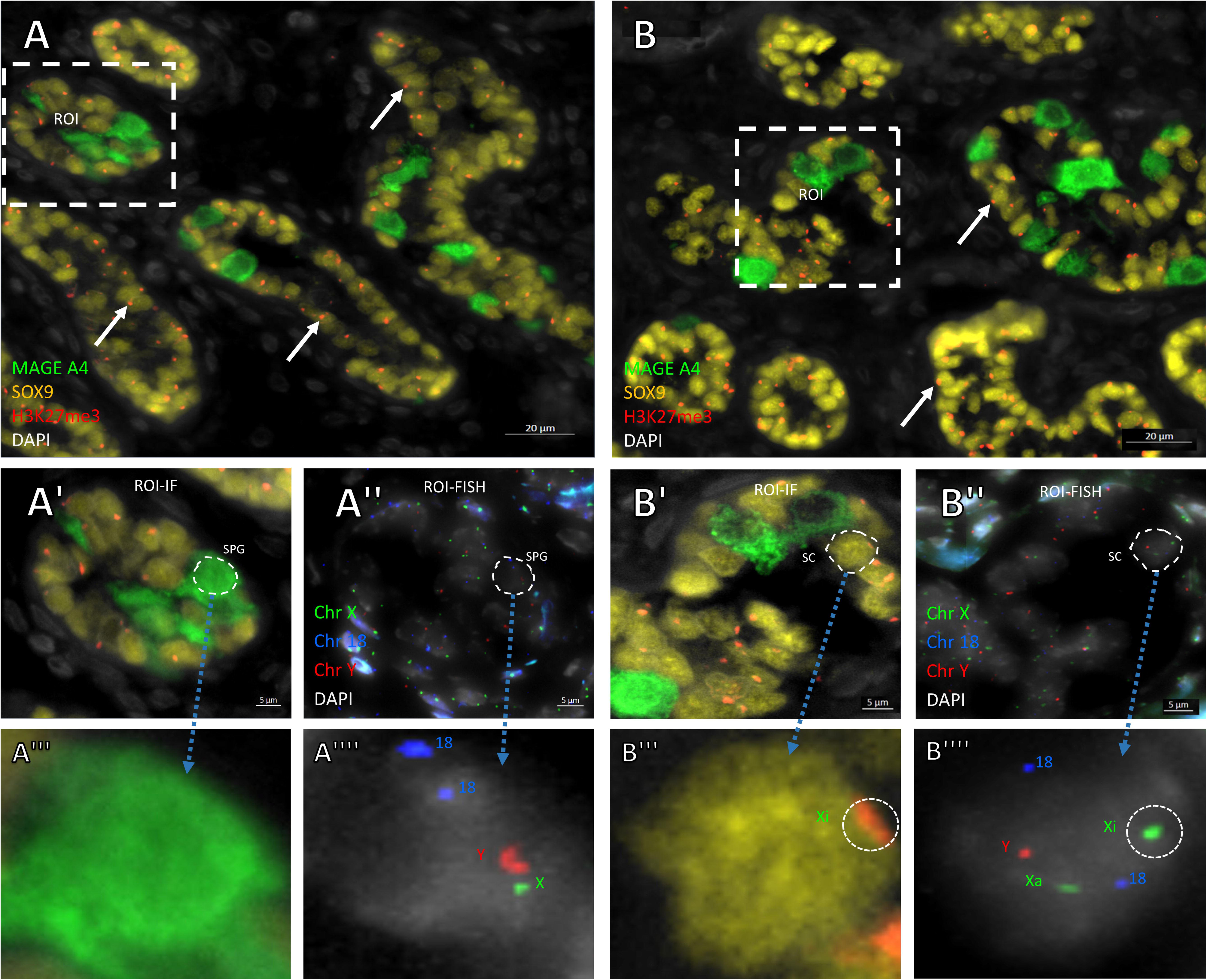
Spermatogonia and Sertoli cells in prepubertal KS patient. **A:** All spermatogonia are Xi^-^. Most Sertoli cells are Xi^+^ in both tubules with or without spermatogonia (arrows). **A’ and A”:** IF and FISH stain of ROI in A, respectively. Spermatogonia are Xi^-^ while almost all Sertoli cells are Xi^+^ (white arrows in A’). **A”’ and A””:** One spermatogonium is magnified. Spermatogonium is negative for Xi (A”’) and has two chr 18, one chr X, and one chr Y signal (A””). **B:** Different area. All spermatogonia are Xi^-^. Most Sertoli cells are Xi^+^ in all tubules (arrows). **B’ and B”:** IF and FISH stain of ROI in B, respectively. Spermatogonia are Xi^-^ while almost all Sertoli cells are Xi^+^ (white arrows in B’). **B”’ and B””:** One Sertoli cell is magnified. It is positive for Xi (encircled Xi in B”’) and has two chr X, two chr 18, and one chr Y signal (B””). Encircled Xi in B”’ overlaps with one (encircled- Xi) of two chr X signals in B””. So, the second green signal is the active X (Xa). **Note:** DAPI was not applied in IF staining because DAPI application prevents the binding of FISH probes, so the pale DAPI colour in IF figures was obtained by autofluorescence of nuclei using the DAPI channel. ROI: region of interest.

**Table 1.** Immunofluorescence analysis for testis specific cells.

|  |  | Tubule analysis |  |  | Spermatogonia |  |  | Sertoli cells |  |  |  |  |  | Peritubular myoid cells |  |  | Leydig cells |  |  |
| --- | --- | --- | --- | --- | --- | --- | --- | --- | --- | --- | --- | --- | --- | --- | --- | --- | --- | --- | --- |
|  |  |  |  |  |  |  |  | Xi<sup>-</sup> tubule |  |  | Xi<sup>+</sup> tubule |  |  |  |  |  |  |  |  |
| Patient |  | # of Xi<sup>-</sup> tubules | # of Xi<sup>+</sup> tubules | # of tubules with SPG | Xi<sup>-</sup> | Xi<sup>+</sup> | % of Xi<sup>+</sup> | Xi<sup>-</sup> | Xi<sup>+</sup> | % of Xi<sup>+</sup> | Xi<sup>-</sup> | Xi<sup>+</sup> | % of Xi<sup>+</sup> | Xi<sup>-</sup> | Xi<sup>+</sup> | % of Xi<sup>+</sup> | Xi<sup>-</sup> | Xi<sup>+</sup> | % of Xi<sup>+</sup> |
| Postpubertal | P1 | 25 | 0 | 1 | 14 | 0 | 0 | 40 | 0 | 0 | - | - | - | 25 | 5 | 17 | 17 | 24 | 59 |
|  | P2 | 53 | 2* | 19 | 381 | 0 | 0 | 25 | 0 | 0 | - | - | - | 20 | 8 | 29 | 18 | 31 | 63 |
|  | P3 | 8 | 0 | 2 | 31 | 0 | 0 | 32 | 0 | 0 | - | - | - | 16 | 2 | 11 | 22 | 38 | 63 |
|  | P4 | 12 | 3 | 3 | 127 | 0 | 0 | 36 | 0 | 0 | 25 | 46 | 65 | 17 | 3 | 15 | 16 | 36 | 69 |
|  | P5 | 3 | 6 | 1 | 46 | 0 | 0 | 23 | 0 | 0 | 11 | 22 | 67 | 31 | 5 | 14 | 17 | 29 | 63 |
|  | P6 | 6 | 0 | 2 | 6 | 0 | 0 | 17 | 0 | 0 | - | - | - | 6 | 2 | 25 | 27 | 42 | 61 |
|  | P7 | 8 | 3 | 8 | 126 | 0 | 0 | 30 | 0 | 0 | 16 | 45 | 74 | 18 | 10 | 36 | 19 | 20 | 51 |
|  | P8 | 3 | 5 | 5 | 93 | 0 | 0 | 16 | 0 | 0 | 11 | 55 | 83 | 15 | 3 | 17 | 9 | 17 | 65 |
|  | P9 | 20 | 1 | 0 | - | - | - | 39 | 0 | 0 | 16 | 27 | 63 | 14 | 5 | 26 | 22 | 30 | 58 |
|  | P10 | 14 | 0 | 0 | - | - | - | 28 | 0 | 0 | - | - | - | 14 | 4 | 22 | 31 | 30 | 49 |
|  | P11 | 4 | 1 | 0 | - | - | - | 45 | 0 | 0 | 3 | 6 | 67 | 14 | 4 | 22 | 30 | 37 | 55 |
|  | P12 | 4 | 0 | 0 | - | - | - | 55 | 0 | 0 | - | - | - | 14 | 6 | 30 | 24 | 34 | 59 |
|  | P13 | 12 | 1 | 0 | - | - | - | 40 | 0 | 0 | 7 | 21 | 75 | 21 | 7 | 25 | 18 | 36 | 67 |
|  | P14 | 12 | 7 | 0 | - | - | - | 35 | 0 | 0 | 8 | 22 | 73 | 9 | 3 | 25 | 18 | 39 | 68 |
|  | P15 | 25 | 10 | 0 | - | - | - | 42 | 0 | 0 | 11 | 19 | 63 | 20 | 4 | 17 | 21 | 11 | 34 |
|  | P16 | 3 | 9 | 0 | - | - | - | 27 | 0 | 0 | 10 | 14 | 58 | 10 | 6 | 38 | 5 | 4 | 44 |
|  | P17 | 0 | 10 | 0 | - | - | - | - | - | - | 9 | 10 | 53 | 8 | 5 | 38 | 14 | 15 | 52 |
| Total |  | 212 | 56 | 41 | 826 | 0 | - | - | - | - | - | - | - | - | - | - | - | - | - |
| Mean±SD |  | - | - | - | - | - | 0±0% | 33±10 | 0±0 | 0±0% | 12±6 | 26±15 | 67±8% | 16±6 | 5±2 | 24±8% | 19±7 | 27±11 | 58±9% |
| Prepubertal | P18 | 0 | 246 | 175 | 300 | 0 | 0 | - | - | - | 4 | 26 | 87 | 11 | 7 | 39 | - | - | - |
|  | P19 | 0 | 690 | 1 | 3 | 0 | 0 | - | - | - | 4 | 26 | 87 | 6 | 6 | 50 | - | - | - |
|  | P20 | 0 | 361 | 6 | 24 | 0 | 0 | - | - | - | 3 | 27 | 90 | 4 | 6 | 60 | - | - | - |
|  | P21 | 0 | - | 0 | - | - | - | - | - | - | 9 | 45 | 83 | 7 | 5 | 42 | 5 | 5 | 50 |
|  | P22 | 0 | - | 0 | - | - | - | - | - | - | 4 | 37 | 90 | 6 | 3 | 33 | - | - | - |
| Total |  | 0 | - | 182 | 563 | 0 | - | - | - | - | - | - | - | - | - | - | - | - | - |
| Mean±SD |  | - | - | - | - | - | 0±0% | - | - | - | 5±2 | 32±8 | 87±3% | 7±2 | 5±1 | 45±10% | - | - | - |

- : not applicable or does not exist. The total count in samples was given for tubules and spermatogonia. Mean±SD per Xi<sup>-</sup> or Xi<sup>+</sup> tubule was given for Sertoli cells, and PMTCs. Mean±SD per randomly selected area was given for Leydig cells. The percentage of Xi<sup>+</sup> cells was given for each cell type.
\* Stands for two exceptional Xi<sup>+</sup> tubules with spermatogonia in P2.

### Sertoli cell analysis

Some tubules of postpubertal samples contained only Xi^-^ Sertoli. These tubules were classified as Xi^-^ tubules, regardless of the presence of spermatogonia. Hence, the percentage of Xi^+^ Sertoli cells in these tubules was 0% (Table 1, Figure 2). In Sertoli cells of Xi^-^ tubules, FISH confirmed that, although there were two signals for chromosome 18, only one signal was found for X and Y chromosomes (Figure 2). The incidence of X and Y chromosome signals of Sertoli cells in Xi^-^ tubules were similar, while more chromosome 18 signals were found (p<0.001) (Figure 2). Therefore, both immunofluorescence and FISH results show that, in postpubertal samples, the karyotype of all Sertoli cells in Xi^-^ tubules is 46,XY, regardless of the presence or absence of spermatogonia.

The mean percentage of Xi^+^ Sertoli cells in Xi^+^ tubules was 67±8% (Table 1, Figure 2). According to the FISH results, most Sertoli cells in Xi^+^ tubules contain two signals for chromosome 18 and X, while only one signal for Y chromosome (Figure 2). In addition, the incidence of chromosome 18 and X signals in these cells was found to be similar, and higher than the Y chromosome (p<0.001), indicating that there are two X chromosomes in these cells (Figure 2). Therefore, it was revealed by both immunofluorescence and FISH that the karyotype of the majority of Sertoli cells in Xi^+^ tubules are 47,XXY.

In prepubertal samples, all tubules with or without spermatogonia were Xi^+^ tubules and the mean percentage of Xi^+^ Sertoli cells was 87±8% in these tubules (Table 1, Figure3). Therefore, the dimorphism for Sertoli cells in terms of Xi positivity which was seen in postpubertal seminiferous tubules was not observed in prepubertal samples. When the FISH results were analyzed, it was observed that Sertoli cells had two signals for both chromosome 18 and X but had only one signal for chromosome Y (Figure 3). Therefore, in prepubertal samples, it was revealed by both immunofluorescence and FISH that the karyotype of all Sertoli cells is 47,XXY, regardless of the presence or absence of spermatogonia.

### Peritubular myoid cell and Leydig cell analysis

Both Xi^+^ and Xi^-^ PTMCs were found around both Xi^+^ and Xi^-^ tubules. (Supplementary figure 2). The rate of Xi^+^ PTMCs was found to be 24±8% in postpubertal and 45±10% in prepubertal samples (Table 1).

In postpubertal samples, the Xi^+^ rate of Leydig cells was found to be 58±9% (Table 1). Leydig cells close to tubules with spermatogonia and those close to Sertoli cell-only tubules with Xi^-^ Sertoli cells were also observed to be both Xi^+^ and Xi^-^ (Supplementary figure 2).

## DISCUSSION

The fact that KS individuals with focal spermatogenesis may have offspring with a 46,XY karyotype thanks to TESE and ICSI (15) shows that euploid spermatozoa are formed in their seminiferous tubules, which raises the question of what is the karyotype of spermatogonia and testis-specific somatic cells in regions where focal spermatogenesis is observed. Our study is the first to reveal the answer to this clinically and scientifically important question both *in situ* and at a cell-specific level.

To date, there have been many publications with conflicting results on the karyotype ofgerm cells that originate focal spermatogenesis (35–40). However, it is generally accepted that only germ cells with the 46,XY karyotype can originate focal spermatogenesis (14, 18, 19). One of the most recent studies on this subject was published by Sciurano et al. The authors claim that while the spermatogonia in focal spermatogenic regions have the 46,XY karyotype, Sertoli cells supporting spermatogonia have the 47,XXY karyotype (14). However, since spermatogonia and Sertoli cells were not evaluated *in situ*, it is not known whether the Sertoli cells belonged to the germ cell-containing tubules or the Sertoli cell-only tubules. In another study conducted by Yamamoto et al., cells were identified according to their shape, volume and position in the tubule, instead of using a cell-specific marker as we did. They reported that spermatogonia were mosaic and all Sertoli cells had the 47,XXY karyotype (37). However, our results suggest that there is no mosaicism amongst spermatogonia as all spermatogonia in KS have the 46,XY karyotype. We demonstrated *in situ* with cell-specific markers that in postpubertal samples, both spermatogonia and Sertoli cells in spermatogonia-containing seminiferous tubules have the 46,XY karyotype, regardless of whether there is ongoing spermatogenesis or not. We detected only two tubules with 46,XY spermatogonia and 47,XXY Sertoli cells in one of the postpubertal patients (supplementary figure 1). However, there was no ongoing spermatogenesis in these tubules. Therefore, in addition to the germ cells that give rise to focal spermatogenesis, Sertoli cells that provide support to these cells also seem to need a 46,XY karyotype, contrary to what was claimed before (14, 37).

It was first demonstrated by Froland et al. that there is dimorphism for the Barr body i.e., Xi, in the seminiferous tubules in KS patients (41), independent of the presence of germ cells. In this study, using traditional histochemical staining methods, it was reported that while Sertoli cells in some tubules were completely devoid of Xi, in other tubules an average of 30% of Sertoli cells was positive for Xi. In our study, we performed high-sensitivity Xi analysis using the H3K27me3 marker (28) in combination with cell-specific markers. Contrary to Froland et al., the average percentage of Xi^+^ Sertoli cells in Xi^+^ tubules in postpubertal samples was found to be 67% in our study. This rate was found to be 87% in prepubertal KS patients.

Fraccaro and Lindsten, who cultured different organs from human fetuses, reported that the incidence of Xi in cultures taken from female fetuses ranged from 34% to 88% (42). In other words, although all cells in females have two X chromosomes, the incidence of Xi varies according to the cell type. In addition, their data are based on cell culture, that is, analysis of structurally intact cells. However, in tissue sections, the incidence of Xi is likely to be lower due to section thickness limitation. As a matter of fact, the incidence of Xi was found to be 31% in female colon muscle cells used as positive control for Xi in our study. Therefore, according to these data, we hypothesize that all Sertoli cells (87% Xi incidence) in the seminiferous tubules in prepubertal individuals and all Sertoli cells (67% Xi incidence) in Xi^+^ tubules in postpubertal individuals have the 47,XXY karyotype. Moreover, PTMCs surrounding both Xi^+^ tubules and Xi^-^ tubules probably have the 47,XXY karyotype (24% and 45% Xi incidence in postpubertal and prepubertal patients, respectively). Similarly, Leydig cells (58% Xi incidence) near and far from the focal spermatogenic regions in postpubertal samples may have the 47,XXY karyotype.

Another controversial issue in the literature is whether germ cells and Sertoli cells lose the extra X chromosome or whether the extra X chromosome cannot be detected because it takes on a different epigenetic structure or gets activated. Froland et al. sees X chromosome loss as unlikely and suggests that two X chromosomes are isopyknotic. However, Sciurano et al., hypothesize that in puberty, Sertoli cells induce spermatogonia differentiation, and during this period some of the spermatogonia lose an X chromosome and survive (14). In contrast, we have shown that spermatogonia have the 46,XY karyotype not only in postpubertal KS but also in prepubertal KS patients of different ages (Table 1). This strengthens the possibility that germ cells lose their extra X chromosome during the fetal period or early post-natal period. The fact that all spermatogonia, without exception, were Xi^-^ in the prepubertal patients we examined, suggests that germ cells that lose their extra X chromosome in the fetal or early postnatal period can survive, whereas germ cells that do not lose their X chromosome cannot. With regard to Sertoli cells, there is no prediction in the literature until now about when these cells lose the extra X chromosome, since it has been thought that Sertoli cells surrounding spermatogonia, have the 47,XXY chromosome pattern. However, as mentioned earlier, we showed that in postpubertal KS samples, besides tubules containing germ cells, many other seminiferous tubules also seem to have Sertoli cells with the 46,XY chromosome pattern. We also showed that in the seminiferous tubules (with or without germ cells) of all prepubertal KS samples, all Sertoli cells have the 47,XXY karyotype. Therefore, we hypothesize that Sertoli cells may also lose one extra X chromosome and this loss occurs during the transition from prepuberty to puberty. However, there are several possibilities that need to be clarified regarding the association of the loss of an extra X chromosome by spermatogonia and Sertoli cells. Since 46,XY spermatogonia are found almost exclusively in Xi^-^ tubules, it is possible that spermatogonia cannot survive if Sertoli cells have not lost an X chromosome. This suggests that spermatogonia can only survive if they have lost an X chromosome and the Sertoli cells in the same tubule also lost an X chromosome. The loss of an X chromosome in both spermatogonia and Sertoli cells can either occur independent or dependent from each other. Sertoli cells may have lost an X chromosome because the germ cells in these tubules have lost an X chromosome, or the other way around. However, as in prepubertal patients, germ cells already had the 46,XY karyotype while Sertoli cells had the 47,XXY karyotype, the former option might be the most likely.

The demonstration that spermatogonia and Sertoli cells might lose one of their X chromosomes raises other questions to be answered: what is the mechanism and exact timing for the extra X chromosome loss in both germ cells and Sertoli cells and, why do Sertoli cells in Xi^+^ tubules not lose the extra X chromosome, while all Sertoli cells in Xi^-^ tubules without exception lose the extra X chromosome?

In conclusion, it was shown for the first time *in situ* with a combination of cell-specific markers that the spermatogonia, whether pre- or postpubertal, have the 46,XY karyotype in non-mosaic KS patients. Therefore, it may be assumed that the spermatogonia that originate focal spermatogenesis have the 46,XY karyotype. In addition to Sertoli cells surrounding spermatogonia in postpubertal samples, Sertoli cells in some tubules, which we classified as Xi^-^ tubules, have the 46,XY karyotype, while Sertoli cells in other tubules, which we classified as Xi^+^ tubules, have the 47,XXY karyotype. In prepubertal patients, Sertoli cells were shown to have the 47,XXY karyotype. Lastly, PTMCs and Leydig cells may have mosaicism in both prepubertal and postpubertal samples, however, further research is needed to confirm these observations.

## Authors contribution statements

S.G. involved in planning the experimental design, data collection, statistical analysis, interpretation of the results, and writing the original draft of the manuscript. V.V. involved in acquisition of postpubertal patient samples and data, review and editing of the manuscript. I.G. involved in acquisition of prepubertal patient samples and data, review and editing of the manuscript. E.G. involved in study conception and design, funding acquisition, supervision of the research, validation of the results, and review and editing of the manuscript.

## Acknowledgments

I sincerely thank Dr. Yves Heremans and Mr. Pierre Hilven for their technical assistance and Dr. Feyza İnceoğlu for her help with statistical analysis.

## Funding

This study was funded by The Scientific and Technological Research Council of Türkiye (TUBİTAK) and a grant (SRP89) from the Vrije Universiteit Brussel.

## Conflict of interest

There is no conflict to declare.

## Figure Caption

**Supplementary figure 1. Two exceptional Xi^+^ tubules with spermatogonia in a postpubertal patient (P2).** Blue arrows show spermatogonia and white arrows show Xi in Sertoli cells. The seminiferous tubule structure is disorganized and does not show a lumen.

**Supplementary figure 2. PTMCs and Leydig cells. A:** PTMCs surrounding a tubule with ongoing spermatogenesis. Blue arrows show Xi^-^ Sertoli cells and yellow arrows show Xi^-^ germ cells. ROI is magnified at the upper right with two Xi^+^ (white arrows) PTMCs. **B:** PTMCs surrounding a SCO tubule with Xi^+^ Sertoli cells. Blue arrows show some of Xi^+^ Sertoli cells. ROI is magnified at the upper right with one Xi^+^ (white arrow) PTMC. **C:** Leydig cells between two tubules with ongoing spermatogenesis. The blue arrow shows Xi^-^ Sertoli cell and the yellow arrows show Xi^-^ germ cells. ROI is magnified at upper right with one Xi^+^ (white arrow) Leydig cell. **D:** Leydig cells from a Leydig cell hyperplastic region. Most of Leydig cells are Xi^+^ (white arrows). ROI is magnified at upper right with one Xi^+^ (white arrow) Leydig cell. **Note:** DAPI was not applied in IF staining because DAPI application prevents the binding of FISH probes, so the pale DAPI colour in IF figures was obtained by autofluorescence of nuclei using the DAPI channel. ROI: region of interest.

---

**Supplementary table 1.** Patients, controls, and testis biopsy sample information.

| Patient |  | Leukocyte Karyotype | Age (year) | Sperm in TESE | Sperm Motility | MAGE A4 | Fixative |
| --- | --- | --- | --- | --- | --- | --- | --- |
| Postpubertal | P1 | 47,XXY | 32 | Yes | Motile | **Positive** | PFA |
|  | P2 | 47,XXY | 31 | Yes | Motile | **Positive** | PFA |
|  | P3 | 47,XXY | 30 | Yes | Immotile | **Positive** | AFA |
|  | P4 | 47,XXY | 27 | Yes | Motile | **Positive** | PFA |
|  | P5 | 47,XXY | 27 | No | - | **Positive** | PFA |
|  | P6 | 47,XXY | 27 | No | - | **Positive** | AFA |
|  | P7 | 47,XXY | 24 | Yes | Immotile | **Positive** | AFA |
|  | P8 | 47,XXY | 14,2 | No | - | **Positive** | AFA |
|  | P9 | 47,XXY | 39 | Yes | Motile | Negative | AFA |
|  | P10 | 47,XXY | 38 | Yes | Motile | Negative | PFA |
|  | P11 | 47,XXY | 38 | No | - | Negative | AFA |
|  | P12 | 47,XXY | 35 | No | - | Negative | AFA |
|  | P13 | 47,XXY | 29 | Yes | Immotile | Negative | AFA |
|  | P14 | 47,XXY | 24 | Yes | Motile | Negative | AFA |
|  | P15 | 47,XXY | 23 | Yes | Motile | Negative | PFA |
|  | P16 | 47,XXY | 15 | No | - | Negative | AFA |
|  | P17 | 47,XXY | 15 | No | - | Negative | AFA |
| Prepubertal | P18 | 47,XXY | 11 | Not applicable | - | **Positive** | AFA |
|  | P19 | 47,XXY | 6 | Not applicable | - | **Positive** | AFA |
|  | P20 | 47,XXY | 4 | Not applicable | - | **Positive** | AFA |
|  | P21 | 47,XXY | 12 | No | - | Negative | AFA |
|  | P22 | 47,XXY | 5 | Not applicable | - | Negative | AFA |
| XX Control |  | 46,XX | 90 | Not applicable | Not applicable | Not applicable | PFA |
| XY Control |  | 46,XY | 47 | Yes | Motile | **Positive** | PFA |

**Supplementary file 1. Immunoflurescence and FISH protocols**

1. IF protocol for H3K27ME3+MAGE A4+SOX9 or H3K27ME3+CYP17A1+ACTA2

1. Xylene, three washes, 5 minutes each.

2. 100% ethanol, two washes, 5 minutes each.

3. 95% ethanol, one wash, 5 minutes.

4. 70% ethanol, one wash, 5 minutes.

5. 50% ethanol, one wash, 5 minutes.

6. dwater 5 minutes.

7. Antigen retrieval: 30 min incubation at 95°C in sodium citrate buffer (pH 6.0).

8. Cool to room temperature for 30 minutes.

9. dwater, one wash, 5 minutes.

10. Draw a circle on the slide around the tissue with a hydrophobic barrier pen.

11. Permeabilization: wash the sections once for 10 minutes with permeabilization buffer containing 1% normal donkey serum and 0.4% Triton X-100 in PBS.

12. Wash in PBS for 10 minutes

13. Blocking: 5% normal donkey serum in PBS-T for 30 minutes at room temperature.

14. Incubate primary antibodies at 4°C overnight.

15. Wash sections twice with 1% serum PBS-T for 10 minutes each.

16. Add a fluorescent label conjugated secondary antibody diluted with 1% serum in PBS and incubate at room temperature for 1 hour.

17. Wash sections once with 1% serum PBS-T for 10 minutes.

18. Wash sections twice with PBS for 10 minutes each.

19. Add Hoechst or DAPI for 5 minutes. (Skipped!)

20. Wash in PBS for 5 minutes. (Skipped!)

21. Coverslip with an anti-fade mounting medium.

2. FISH protocol

Day 1

Preparatory steps

1. Prepare two ethanol series (70%, 90%, and 100% ethanol solutions): Dilute 100% ethanol with deionized or distilled water.

2. Heat Pretreatment Solution Citric (PT1): Warm to 98°C.

3. Wash Buffer SSC (WB1): Bring to room temperature (RT).

4. ZytoLight FISH Probe: Bring to RT before use, and protect from light.

Pretreatment

(After the scanning of immunofluorescence stained slides)

1. Remove the coverslips by immersing the slides in PBS for about 5 minutes.

4. Wash slides 2x 2 min in deionized water.

5. Incubate for 15 min in pre-warmed Heat Pretreatment Solution Citric (PT1) at 98°C.

6. Transfer slides immediately to deionized or distilled water, wash for 2x 2 min, and drain off or blot off the water.

7. Apply (dropwise) Pepsin Solution (ES1) to the specimens and incubate for 15 min at 37°C in a humidity chamber.

8. Wash for 5 min in Wash Buffer SSC (WB1).

9. Wash for 1 min in deionized or distilled water

10. Dehydration: in 70%, 90%, and 100% ethanol, each for 1 min

11. Air dry sections.

Denaturation and hybridization

1. Pipette 10 μl of the ZytoLight FISH Probe onto each pretreated specimen.

2. Cover specimens with a coverslip and seal the coverslip.

3. Place slides on a hot plate or hybridizer and denature specimens for 10 min at 75°C.

4. Transfer the slides to a humidity chamber and hybridize overnight at 37°C.

Day 2 Preparatory steps

1. Preparation of 1x Wash Buffer A: Dilute 1 part 25x Wash Buffer A (WB2) with 24 parts deionized or distilled water. Fill three staining jars with the 1x Wash Buffer A and pre-warm it to 37°C.

2. DAPI/DuraTect-Solution (MT7): Bring to room temperature before use, protect from light.

Post-hybridization and detection

1. Carefully remove the rubber cement or glue.

2. Remove the coverslip by submerging in 1x Wash Buffer A at 37°C for 1-3 min.

3. Wash using 1x Wash Buffer A for 2x 5 min at 37°C.

4. Incubate the slides in 70%, 90%, and 100% ethanol, each for 1 min.

5. Air dry the samples protected from light.

6. Pipette 25 μl DAPI/DuraTect-Solution (MT7) onto the slides. Cover the samples with a coverslip. Incubate in the dark for 15 min.

**Supplementary table 2.** Primary and secondary antibody information.

| **Antibody** | **Product code** | **Dilution** | **Provider** |
| --- | --- | --- | --- |
| Rabbit anti-MAGE-A4 | 82491 | 1/200 | Cell Signaling Technology, Leiden, Netherlands |
| Goat anti-SOX9 | AF3075 | 1/200 | BioTechne, Minneapolis, USA |
| Rabbit anti-CYP17A1 | 14447-1-Ap | 1/300 | Proteintech, Manchester, UK |
| Goat anti-ACTA2 | NB300-978 | 1/100 | Novusbio, Abingdon, UK |
| Mouse anti-H3K27me3 | ab6002 | 1/200 | Abcam, Cambridge, UK |
| Donkey anti-rabbit Alexa Flour 488 | A21202 | 1/200 | Life Technologies, California, USA |
| Donkey anti-goat Alexa Flour 647 | A21447 | 1/200 | Life Technologies, California, USA |
| Donkey anti-mouse Alexa Flour-594 | A21203 | 1/200 | Life Technologies, California, USA |

## Notes

### Competing Interest Statement

The authors have declared no competing interest.

