## Supplementary figures and images for "Testicular mosaicism in non-mosaic postpubertal Klinefelter patients with focal spermatogenesis and in non-mosaic prepubertal Klinefelter boys"

### Supplementary figure 1

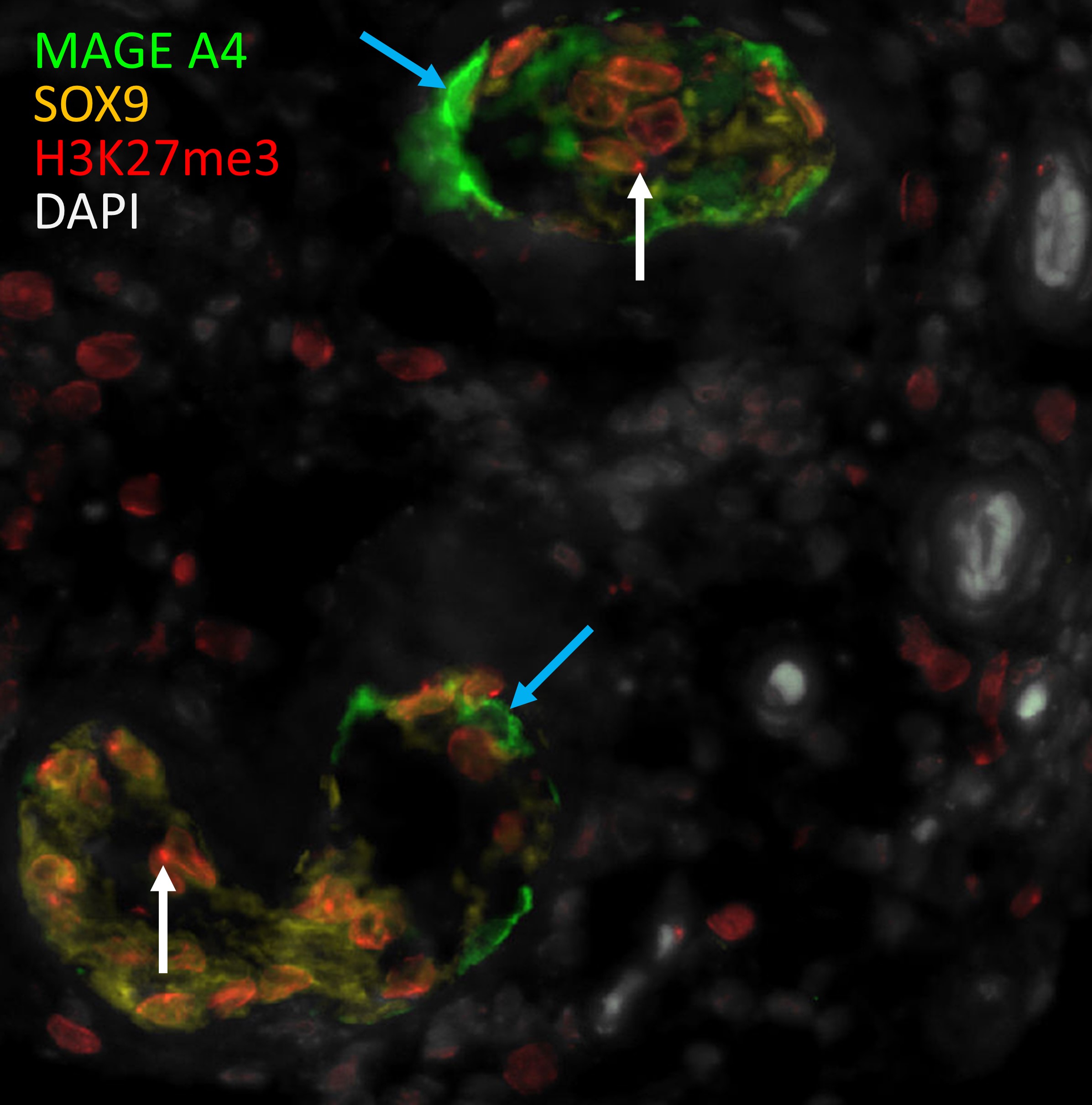

### Supplementary figure 2

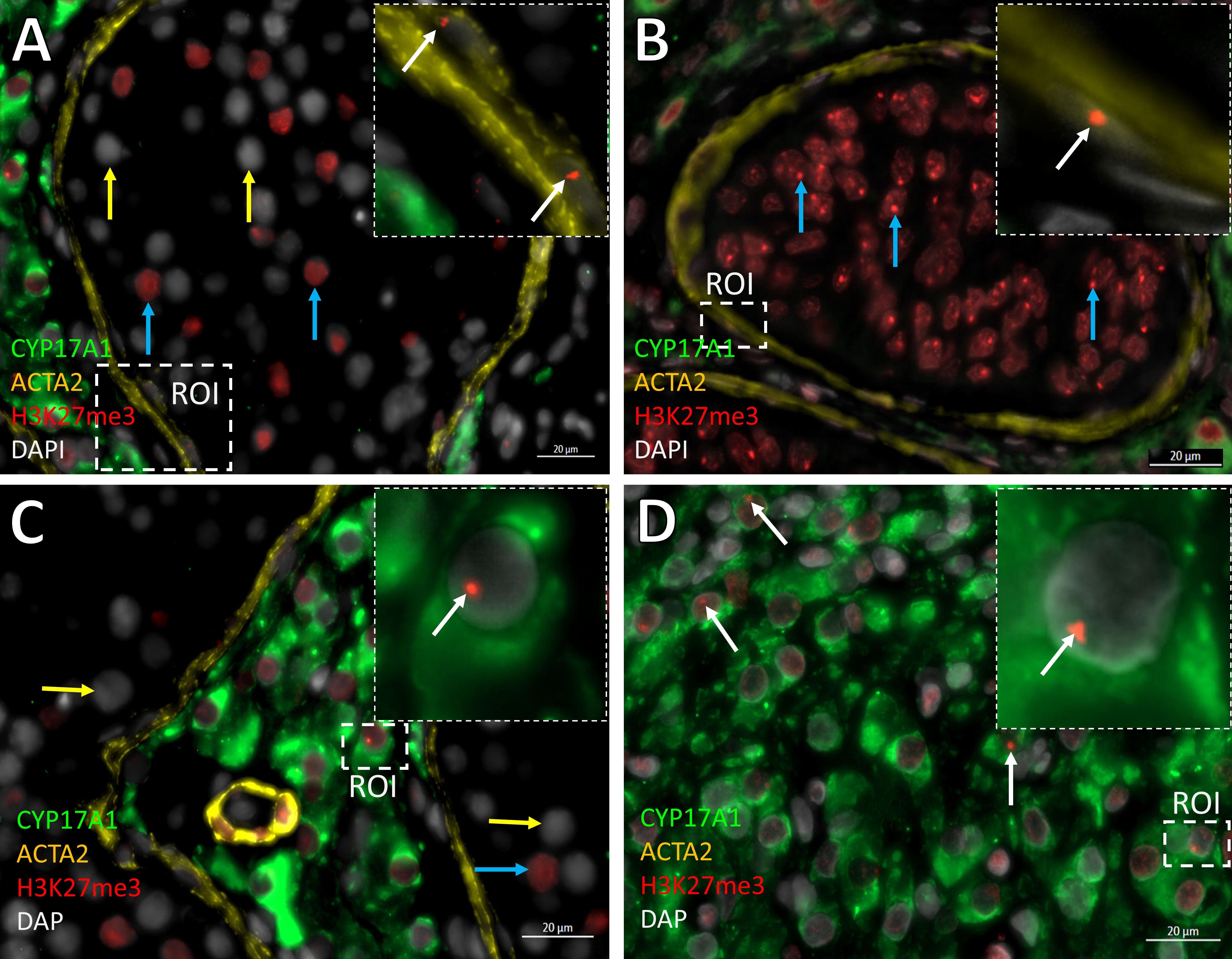
